# Mouse Adapted SARS-CoV-2 protects animals from lethal SARS-CoV challenge

**DOI:** 10.1101/2021.05.03.442357

**Authors:** Antonio Muruato, Michelle N. Vu, Bryan A. Johnson, Meredith E. Davis-Gardner, Abigail Vanderheiden, Kumari Lokugmage, Craig Schindewolf, Patricia A. Crocquet-Valdes, Rose M. Langsjoen, Jessica A. Plante, Kenneth S. Plante, Scott C. Weaver, Kari Debbink, Andrew L. Routh, David Walker, Mehul S. Suthar, Xuping Xie, Pei-Yong Shi, Vineet D. Menachery

## Abstract

The emergence of SARS-CoV-2 has resulted in a worldwide pandemic causing significant damage to public health and the economy. Efforts to understand the mechanisms of COVID-19 disease have been hampered by the lack of robust mouse models. To overcome this barrier, we utilized a reverse genetic system to generate a mouse-adapted strain of SARS-CoV-2. Incorporating key mutations found in SARSCoV-2 variants, this model recapitulates critical elements of human infection including viral replication in the lung, immune cell infiltration, and significant *in vivo* disease. Importantly, mouse-adaptation of SARS-CoV-2 does not impair replication in human airway cells and maintains antigenicity similar to human SARS-CoV-2 strains. Utilizing this model, we demonstrate that SARS-CoV-2 infected mice are protected from lethal challenge with the original SARS-CoV, suggesting immunity from heterologous CoV strains. Together, the results highlight the utility of this mouse model for further study of SARS-CoV-2 infection and disease.

## Introduction

Severe acute respiratory syndrome coronavirus 2 (SARS-CoV-2), the virus that causes COVID-19 disease, emerged in late 2019 and has since caused an ongoing outbreak with over 153 million cases and over 3.2 million deaths in the last 17 months^1,2^. The novel coronavirus, similar to previous emergent SARS-CoV and Middle East Respiratory Syndrome (MERS)-CoV, can produce severe respiratory disease characterized by fever, labored breathing, pulmonary infiltration and inflammation^3,4^. In severe cases, SARS-CoV-2 can lead to acute respiratory distress and death. Unlike the earlier pandemic CoVs, SARS-CoV-2 maintains the ability to spread asymptomatically and causes a range of disease from mild to severe^5^. These factors have led to a world-wide outbreak that continues to rage over a year after its emergence.

In responding to the outbreak, understanding the complexity of SARS-CoV-2 infection has been hampered by the limitations of small animal models ^6^. Early on, wild-type SARS-CoV-2 was shown to be unable to utilize mouse ACE2 for entry and infection ^7^. Alternative models utilized receptor transgenic mice expressing human ACE2 or Syrian golden hamsters to evaluate SARS-CoV-2 infection and disease *in vivo* ^6^. However, the transgenic models, while causing severe disease and lethality, have distinct infection tropism leading to encephalitis in addition to lung disease ^8-10^. Similarly, while the hamster model has provided utility in studying disease and transmission ^11^, the absence of genetic knockout and immunological tools limits the types of studies that can be pursued. Without a robust mouse model, many of the resources used to study infection and the immune response are unavailable for SARS-CoV-2 experiments.

In order to alleviate these issues, we set out to develop a mouse-adapted strain of SARS-CoV-2 using standard laboratory strains. Building from our infectious clone system ^12^, we incorporated amino acid changes that facilitated replication in standard Balb/C mice and serially passaged the mutant to create a mouse-adapted strain (CMA3p20) that causes significant weight loss, disease, and lung damage following infection. Notably, virus replication in this model is limited to the respiratory system, thus recapitulating disease observed in most humans. Importantly, the SARS-CoV-2 CMA3p20 strain did not attenuate replication in primary human airway cultures or change the antigenicity of the mouse-adapted strain relative to WT control virus, making it suitable for vaccine and therapeutic studies. Finally, following prior infection with SARS-CoV-2 CMA3p20, mice were protected from lethal challenge with SARS-CoV despite the absence of sterilizing immunity. Together, the results highlight the utility of SARS-CoV-2 CMA3p20 to study infection and pathogenesis in standard mouse lines.

## Results

The initial, emergent strains of SARS-CoV-2 had spike proteins unable to utilize mouse ACE2 and infect standard laboratory mice ^7^. To overcome this barrier, we generated a series of mutations in the receptor-binding domain (RBD) of SARS-CoV-2 using our infectious clone ^12^. Our initial efforts modeled the interaction between SARS-CoV-2 and mouse ACE2 and used previous mouse adapted strains of SARS-CoV (MA15, MA20, and v2163) ^13^ to design mutants including changes at Y449H (MA1), Y449H/L455F (MA2), and F486L/Q498Y (MA4) (**S. Fig. 1A-C**). We also generated a series of mutants based on a reported natural SARS-CoV-2 isolate (MASCP6) capable of infecting mice^14^, which has spike change at N501Y and several additional mutations (**S. Fig. 2A**). Given the capacity of the MASCP6 strain to replicate in mice, we generated mutants that had the spike mutation alone (CMA1), the spike/N protein mutation (CMA2), and all four changes (CMA3) (**S. Fig. 2A**). For each of the six mutants, we utilized site-directed mutagenesis in the WA1 strain clone and rescued virus stocks on Vero E6 cells (**S. Fig. 2B**). We subsequently infected 10-week-old female Balb/C mice with 10^5^ plaque forming units (PFU) of each mutant virus and evaluated replication in the lung 2 days post infection. For WT, MA1, and MA2, no evidence of viable infection was detected in mouse lung tissues (**S. Fig. 2C**); however, MA4 and CMA1-3 had robust replication in mouse lung suggesting multiple combinations of RBD changes could provide compatibly with mouse ACE2 sufficient for replication in a standard laboratory mouse strain.

To further evaluate the mouse adapted strains, we focused on SARS-CoV-2 CMA1, CMA2, and CMA3 mutants over a four-day time course. In female ten-week-old Balb/c mice infected with 10^5^ PFU, none of three mutants induced major disease (**S. Fig. 3A**), although both CMA2 and CMA3 caused more weight loss than CMA1. Examining viral replication in the lung, all three mutants produced ~10^5^ PFU per lobe at day 2 post infection (**S. Fig. 3B**). However, no virus was detected at day 4, suggesting rapid clearance by the host. To determine if type I interferon was the major factor blunting infection, IFNAR^-/-^ SJV129 mice were infected with CMA1, CMA2, and CMA3 at 10^5^ PFU. Following infection, all three CMA mutant strains caused significant disease with both CMA2 and CMA3 peaking at ~10% weight loss (**S. Fig. 3C**). However, despite increased disease, viral titers were only slightly higher at day 2 than immune competent Balb/c mice and still cleared by day 4 for all three strains (**S. Fig. 3D**). Together, the results indicate that SARS-CoV-2 CMA1, CMA2, and CMA3 can replicate in both Balb/C and IFNAR^-/-^ mice, but fail to sustain continued replication *in vivo*.

### Serial passage of SARS-CoV-2 CMA3

In order to generate a SARS-CoV-2 strain that produced significant disease in an immune competent mouse, we serially passaged SARS-CoV-2 CMA3 in 10-week-old Balb/c mice. A single mouse was infected with 10^5^ PFU of CMA3 (p0); the mouse was subsequently euthanized at 1 day post infection with half the lung lobes taken for viral RNA and the other lobes homogenized, clarified, and used to inoculate subsequent passages (**Fig. 1A**); lung samples were titered by plaque assay to verify continued SARS-CoV-2 replication (**Fig. 1B**). After passages (p) 10, 15, and 20, stock viruses were generated on Vero E6 cells, used to infect 10-week-old Balb/C mice, and compared to the disease caused by the original CMA3 p0 strain (**Fig. 1C**). Following 10^5^ PFU challenge, mice infected with p10 and p15 were found to have augmented weight loss compared to p0; however, mice infected with p20 showed 10% weight loss by day 3 and signs of disease including ruffled fur and hunched posture. We subsequently deep sequenced the passaged virus from the lung RNA and identified two additional spike mutations (K417N and H655Y) and a mutation in the E protein (E8V). Several other mutations were also found as minority variants in the spike and in other parts of the genome. Modeling the receptor binding domain interaction (**Fig. 1D**), K417N and N501Y likely improve binding to mouse ACE2 and facilitate increased *in vivo* disease. Together, mouse adaptation of SARS-CoV-2 CMA3 incorporated three additional fixed mutations that drive increased disease in mice.

**Figure 1.**
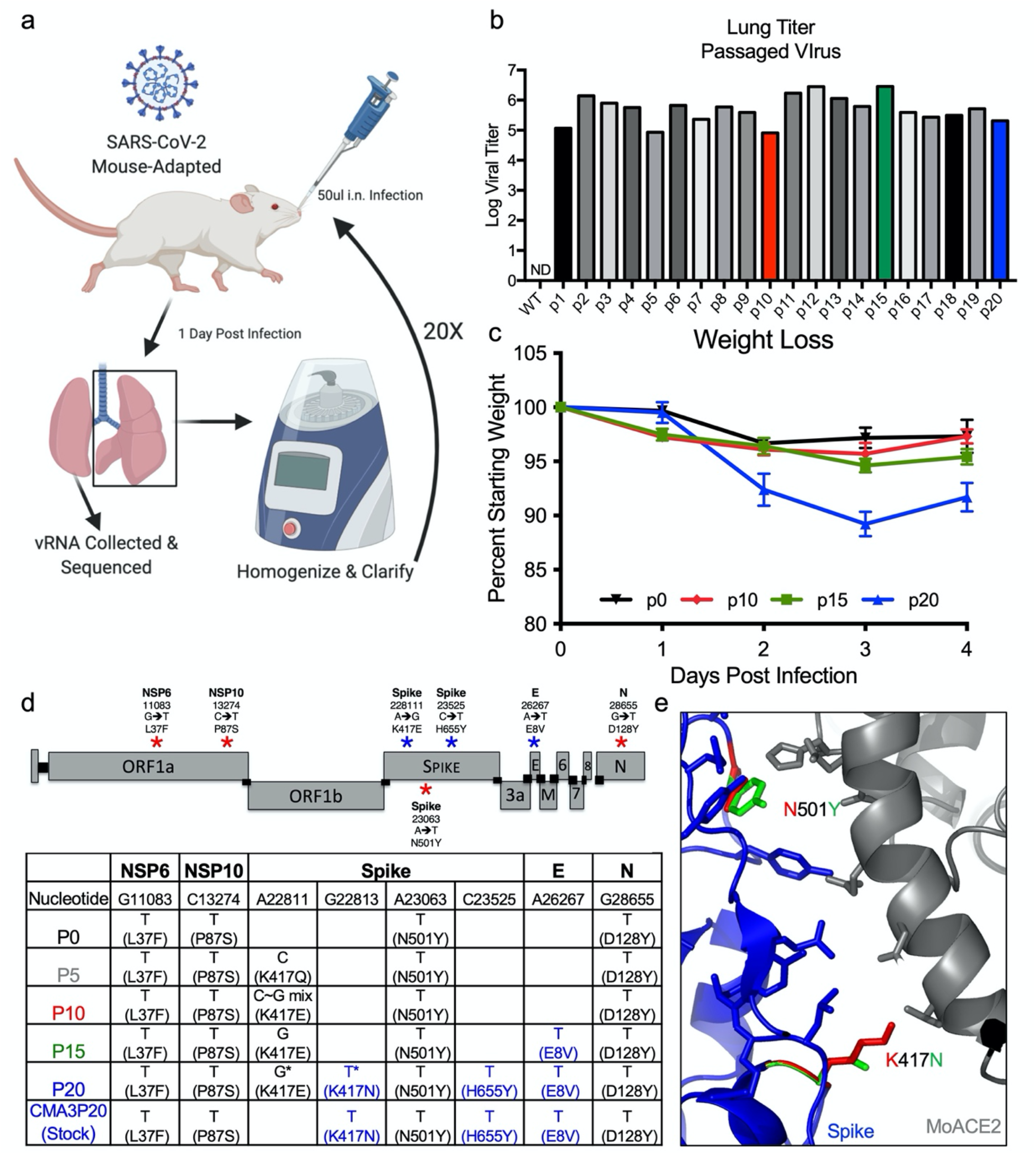
Mouse-adaptation of SARS-CoV-2. a) Schematic of adaptation of SARS-CoV-2 CMA3p20. One ten-week-old female Balb/c mice was infected with SARS-CoV-2 CMA3 for 1 day, euthanized, and lung tissues harvested for viral RNA and viral titer determination. Lung tissues were homogenized, clarified, and 50ul used to inoculate subsequent animals for 20 passages (p). b) Viral replication of CMA3 p1-p20 from lung homogenates isolated from infected mice 1 day post infection. c) Stock virus generated at passages 0, 10, 15, and 20 was used to infect 5 female Balb/c mice at 10^5^ PFU and evaluated for weight loss over a 4-day time course. d). Schematic of engineered (red stars) and passage-acquired (blue stars) mutations in CMA3p20 stock virus. Table includes Sanger equivalent accumulation of mutations over passages p5, p10, p15, p20, and final stock used for subsequent studies. e) Modeling RBD spike mutations N501Y and K417N found in CMA3p20 with mouse ACE2.

### Characterization of CMA3p20

Having observed significant disease in mice infected with CMA3p20 relative to the initial strain of CMA3, we next evaluated weight loss, viral replication, and histopathology in Balb/c mice. First, we tested CMA3p20 for a dose-dependent impact on weight loss (**S. Fig. 4A**); both 10^6^ and 10^5^ PFU caused significant, dose-dependent weight loss with minimal disease observed in the 10^4^ challenge. We also compared CMA3p20 infection associated weight loss to a B.1.1.7 SARS-CoV-2 variant (UK) which contains the N501Y mutation that permits virus replication in mice (**S. Fig. 3A-D**). After challenge with the 10^6^ PFU of the B.1.1.7 variant, female Balb/C mice lost approximately 10% of their starting weight by day 2 and recovered (**S. Fig. 4B**). Together, the results indicate more severe disease with the mouse adapted CMA3p20 than the B.1.1.7 variant.

We subsequently used the 10^5^ PFU dose of CMA3p20 to examine infection compared to SARS-CoV-2 CMA3 over a seven-day time course. Following infection, 10-week-old female Balb/C mice CMA3p20-infected mice lost significant weight over the first four days, peaking at day 3 with >10% weight loss (**Fig. 2A**). In contrast, the original CMA3 caused minimal weight loss over the course of the seven-day infection. We next examined viral replication in the lung at days 2, 4, and 7 post infection (**Fig. 2B**). CMA3p20 infection had a significant 0.5 log increase in viral load over CMA3 in the lung at day 2; this difference was diminished at day 4 (0.25 log increase, not statically significant) and both virus strains were cleared by day 7 in the lung. We also observed day 2 replication in the trachea of mice that was cleared by day 4 in both CMA3 and CMA3p20 infection (**Fig. 2C**). Together, the data demonstrate robust weight loss and clear replication in the mouse respiratory tract.

**Figure 2.**
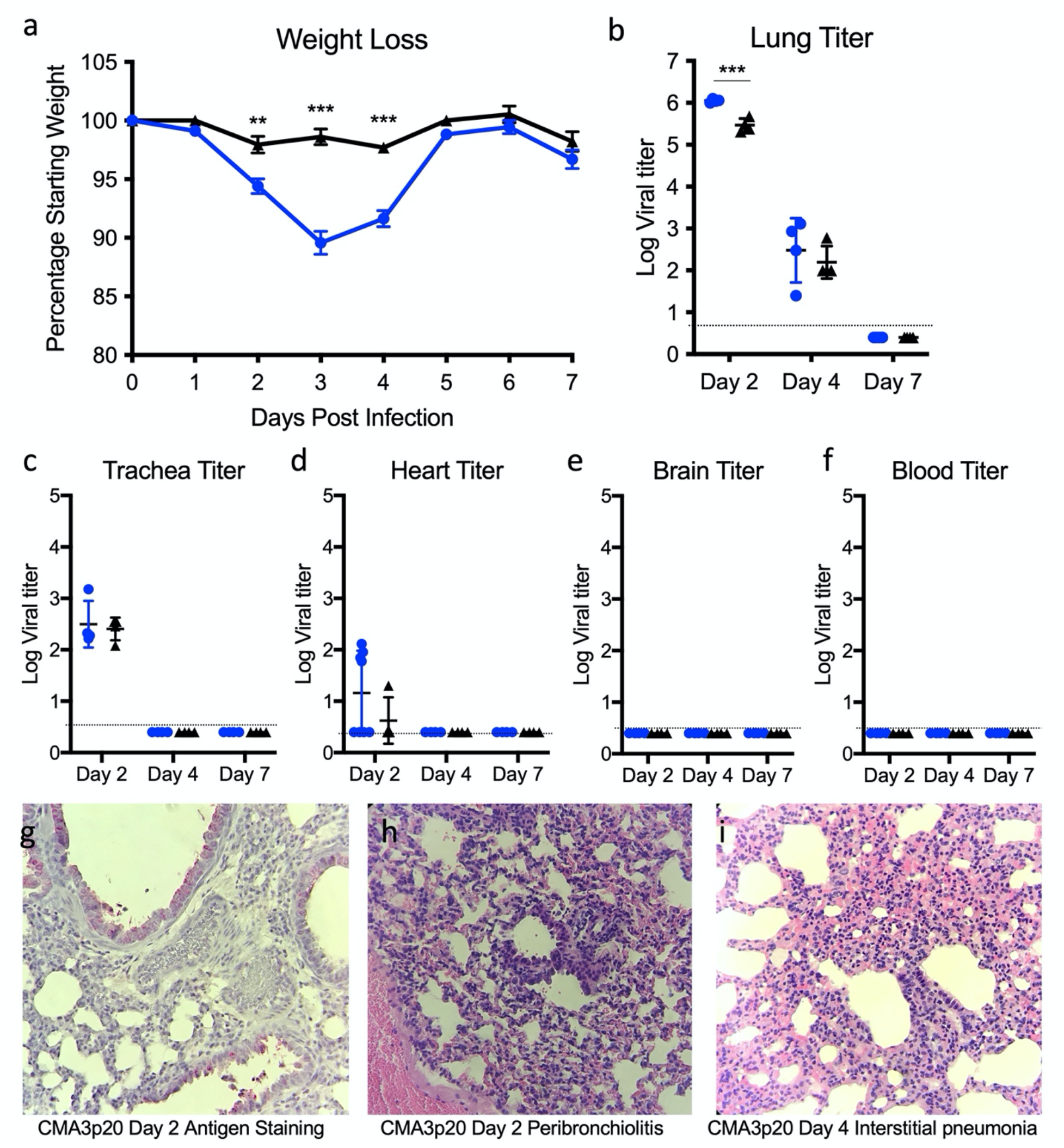
SARS-CoV-2 CMA3p20 induces disease restricted to the lung. a-f) Ten-week-old Balb/c mice were infected with 10^5^ PFU of SARS-CoV-2 CMA3 (black) or CMA3p20 (blue) and followed for a) weight loss and viral titer in the b) lung, c) trachea, d) heart, e) brain, and f) blood. g-i) Histology from CMA3p20 infected mice showed g) viral antigen (N-protein) staining in the airways and parenchyma at day 2. Significant lung infiltration, inflammation and damage was observed at h) day 2 and i) day 4 post infection. Magnification at 10x for g-i.

We next evaluated SARS-CoV-2 replication in non-respiratory tissues. Following infection, we noted replication in the heart tissue of a subset of animals at day 2 (**Fig. 2D**). However, infection was transient and not uniform in all animals and no virus was detected in the heart at later time points. We subsequently evaluated viral load in the brain and blood and found no evidence for CMA3 or CMA3p20 infection by plaque assay (**Fig. 2E-F**). To further verify viral replication, we also examined viral RNA expression in the lung, heart, brain, spleen, and liver (**S. Fig. 4C**). While robust viral RNA was observed in the lung, the other tissues had minimal evidence for CMA3p20 replication. Together, the data indicate that the SARS-CoV-2 CMA3p20 strain is primarily restricted to and disease driven by virus replication in the respiratory tract.

### CMA3p20 induces significant immune infiltration and lung damage

Further examining lung tissue, histopathology analysis of CMA3p20 infection indicated robust virus replication, immune infiltration, and tissue damage. Utilizing antigen staining against the N protein, we saw evidence for viral replication primarily in the bronchioles with additional staining in the lung parenchyma at day 2 post infection (**Fig. 2G**). We also observed lung infiltration and inflammation following CMA3p20 challenge characterized by peribronchioloitis, perivascular cuffing, and perivasculitis by day 2 post infection (**Fig. 2H**). Similarly, at day 4, we noted collapsed airways and interstitial pneumonia (**Fig. 2I**). Some portions of the day 4 lungs infected with CMA3p20 also had virus induced damage including enlarged and multinucleated alveolar type II cells (**S. Fig. 5A**), loss of cellular polarity (**S. Fig. 5B**), and immune cells in the bronchiolar lumen (**S. Fig. 5D**). Together, the histopathology results demonstrated significant damage, inflammation, and disease in the lung following infection with SARS-CoV-2 CMA3p20.

### CMA3p20 retains replication capacity in primary human respiratory cells

Altering SARS-CoV-2 to be permissive in mice can impact its replication capacity in human cells ^15^. Therefore, we examined the ability of CMA3p20 to replicate in primary human airway epithelial (HAE) cultures compared with the WT SARS-CoV-2 WA1 strain. Grown on an air liquid interface, primary HAEs represent a useful *in vitro* model of the human airway ^16^. Following infection, CMA3p20 had equal replication to WT SARS-CoV-2 over a 72-hour time course in primary HAE cultures (**Fig. 3A**). Similarly, viral RNA levels at 72 hours post infection were equivalent between CMA3p20 and SARS-CoV-2 WA1 strain (**Fig. 3B**). Together, the results indicate that mouse adaption resulted in no significant replication attenuation of CMA3p20 in primary human airway cells.

**Figure 3.**
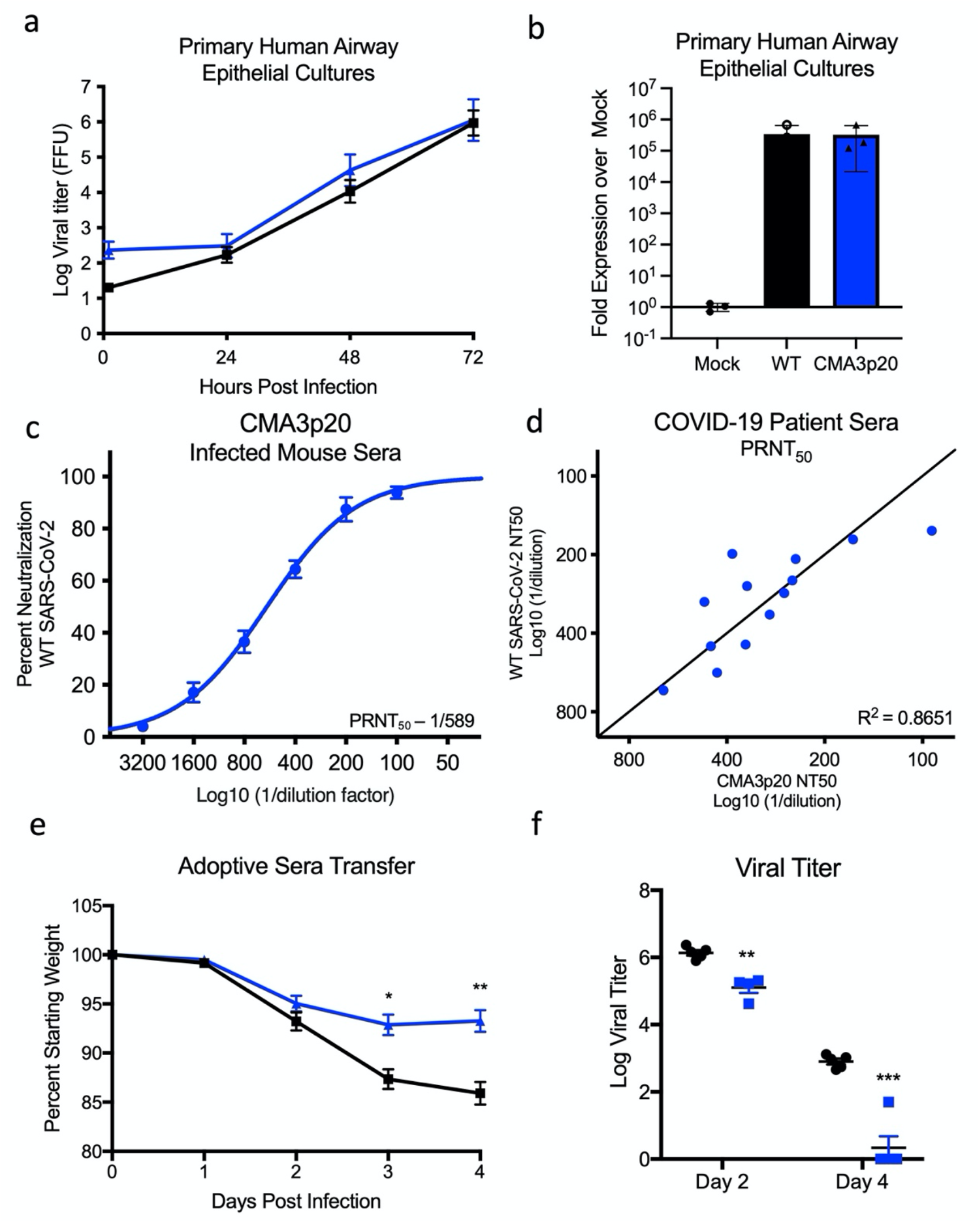
CMA3p20 strain maintains human replication capacity and antigenicity. a-b) Primary human airway cultures were infected with SARS-CoV-2 WT (black) or CMA3p20 (blue) at an MOI of 0.01 and evaluated for a) viral titer and b) viral RNA. c) Sera collected from female Balb/c mice 28 days post infection with 10^6^ PFU of SARS-CoV-2 CMA3p20 were evaluated for capacity to neutralize WT SARS-CoV-2 via PRNT50 assay. d) PRNT50 values from COVID19 patient sera plotted against WT virus (y-axis) versus CMA3p20 virus (x-axis). e-f) Ten-week-old female Balb/c mice were treated intraperitoneally with 100ul of human COVID19 sera or control (PBS) one day prior to infection. Mice were subsequently challenged with 10^5^ PFU of SARS-CoV-2 CMA3p20 and evaluated for e) weight loss and f) viral titer in the lung.

### CMA3p20 retains antigenicity similar to WT SARS-CoV-2

In addition to differences in replication in human cells, spike changes in SARS-CoV-2 could alter the overall antigenicity of CMA3p20 as compared to SARS-CoV-2 derived from humans; this result would make it more difficult to interpret vaccine and protection studies derived from mice. Therefore, to evaluate antigenicity, we infected 10-week-old female Balb/C mice with 10^6^ PFU of CMA3p20, then, euthanized and harvested sera 28 days post infection. We subsequently used the mouse sera to measure plaque reduction neuralization titer (PRNT_50_) against the wild-type SARS-CoV-2 WA1 strain (**Fig. 3C**). Mouse sera from mice (n=7) infected with CMA3p20 neutralized WT SARS-CoV-2 with a PRNT_50_ value of ~1/600. To further evaluate CMA3p20 antigenicity, we examined PRNT_50_ assays utilizing sera from acutely infected COVID-19 patients (**Fig. 3D**). Performing neutralization assays in parallel, we found that CMA3p20 had PRNT^50^ values similar to WT SARS-CoV-2 with each COVID19 patient serum tested. With a R^2^ value of 0.8651 over the 13 samples, the results indicated that CMA3p20 retains similar antigenicity to the WT SARS-CoV-2 and has utility for vaccine and protection studies.

To further demonstrate the utility of CMA3p20 to understand *in vivo* protection, we performed a passive transfer experiment with COVID-19 patient sera. One day prior to infection, 10-week-old female BALB/C mice were pre-treated intraperitoneally with either control (PBS) or 100ul of convalescent serum from a COVID-19 patient. Mice were subsequently challenged with 10^5^ PFU of CMA3p20 and monitored for weight loss and viral titer. Mice treated with acutely infected COVID-19 patient serum had significantly reduced weight loss at day 3 and 4 post infection as compared to control mice (**Fig. 3E**). Similarly, viral titers in the lung were reduced at both day 2 and day 4 in mice receiving COVID-19 patient serum as compared to control. Consistent with the PRNT_50_ data (**Fig. 3D**), the results from the passive transfer experiment demonstrate that antibody-based immunity generated following human infection can effectively neutralize SARS-CoV-2 CMA3p20. Together, the results confirm similar antigenicity of CMA3p20 and WT SARS-CoV-2.

### Prior SARS-CoV-2 infection protects from lethal SARS-CoV challenge

Having established a SARS-CoV-2 mouse model with significant disease, we next evaluated the capacity of CMA3p20 to protect against heterologous SARS-CoV challenge. Ten-week-old female Balb/c mice were infected with 10^6^ PFU of CMA3p20 or control (PBS), monitored for weight loss, and allowed to recover. CMA3p20 infected and control mice were subsequently challenged with a lethal dose of mouse-adapted SARS-CoV (10^4^ PFU) ^17^. Control mice infected with SARS-CoV MA15 had rapid weight loss and lethality with all mice reaching euthanasia criteria by day four post infection (**Fig. 4A & B**). In contrast, mice previously infected with SARS-CoV-2 CMA3p20 had less weight loss compared to controls, only losing approximately 10% of their starting weight by day 2. The CMA3p20-infected mice recovered their starting weight at late times demonstrating protection from lethal SARS-CoV infection. Examining disease score, mice previously infect with SARS-CoV-2 CMA3p20 showed some disease at day 2, but were generally devoid of ruffled fur, diminished movement, or hunching (**Fig. 4C**). In contrast, the mock infected animals challenged with SARS-CoV showed significant disease that escalated over the course of infection and required euthanasia by day 4 for all animals remaining in the study (n=10). For both CMA3p20 infected and uninfected animals, robust SARS-CoV replication was observed in the lung (**Fig. 4D**). However, mice infected with CMA3p20 has a significant reduction in viral loads as compared to control animals. Yet, the viral replication indicates sterilizing immunity was not achieved. We subsequently evaluated the neutralization capacity of SARS-CoV-2 CMA3p20 sera against SARS-CoV (**Fig. 4E**). CMA3p20 mouse sera from pre-challenge, day 2, day 4, and 7 post SARS-CoV infection were able to neutralize SARS-CoV with a low range of PRNT_50_ values (**Fig. 4F**). Both the pre-challenge and day 2 post challenge sera had neutralization levels <1/150, while day 4 and 7 post-challenge sera were only augmented to ~1/200. While significantly less neutralization than what is observed against WT SARS-CoV-2 (~1/600), the results suggest that SARS-CoV-2 CMA3p20 infection induces protection sufficient to protect from lethal challenge with SARS-CoV.

**Figure 4.**
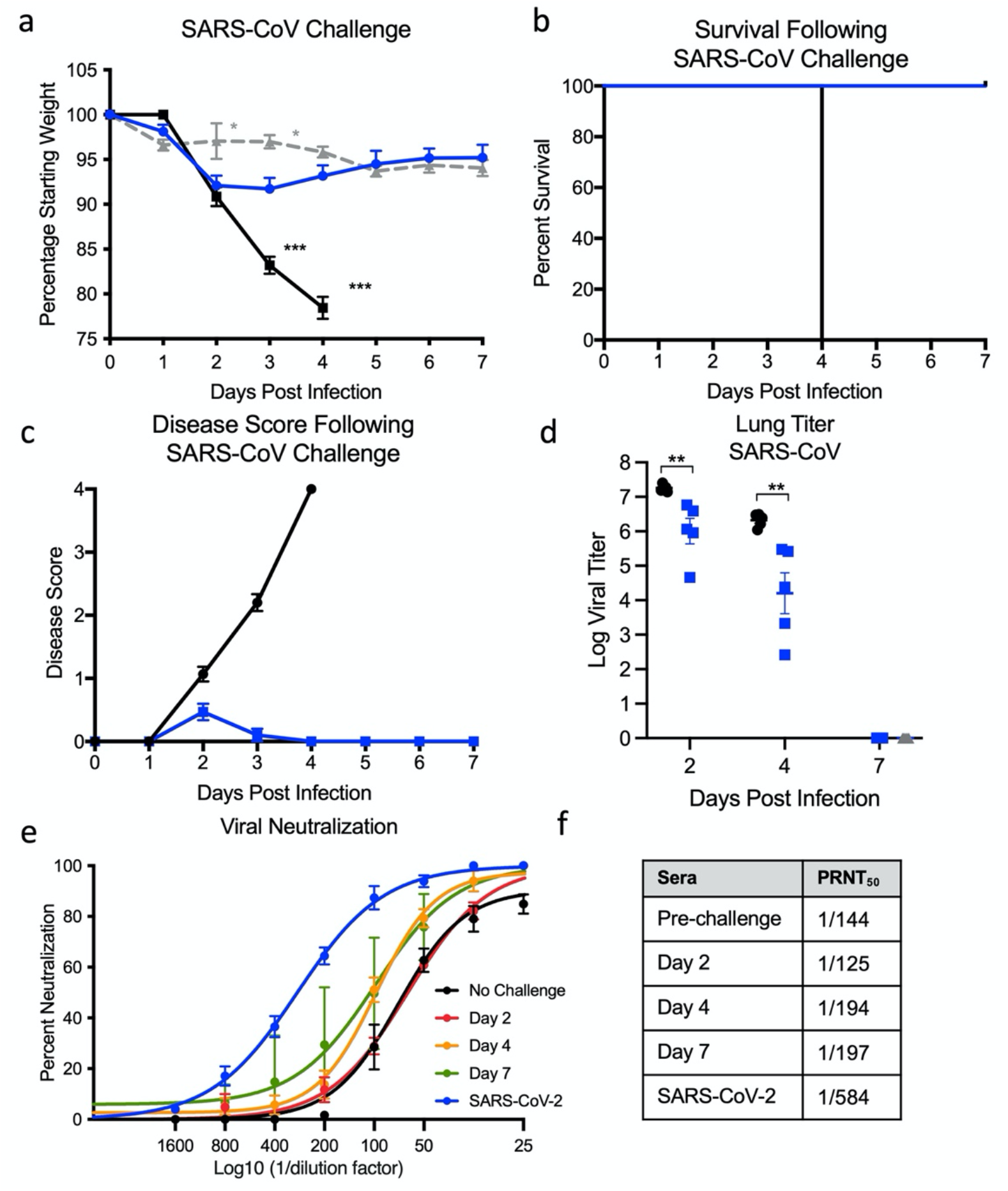
Prior infection with SARS-CoV-2 protects from lethal SARS-CoV challenge. a-c) Ten-week-old female Balb/c mice were previously infected with 10^6^ PFU of SARS-CoV-2 CMA3p20 (blue) or mock (black), monitored for weight loss, and allowed to recover. Twenty-eight days post infection, both groups were challenged with a lethal dose (10^4^ PFU) of mouse-adapted SARS-CoV and evaluated for a) weight loss, b) lethality, and c) disease score. d) Mice were subsequently euthanized at day 2, 4, and 7 and lung tissue examined for viral replication. e) Sera from CMA3p20 infected and SARS-CoV challenged were evaluated for virus neutralization (PRNT50) against SARS-CoV-2 (blue) or SARS-CoV over time (no rechallenge-black, day 2 red, day 4-orange, day 7-green).

## Discussion

In this manuscript, we utilized a reverse genetic system ^18^ and *in vivo* adaptation to generate a mouse-adapted strain of SARS-CoV-2 (CMA3p20). CMA3p20 induces a dose-dependent disease in young, female Balb/C mice with viral replication limited primarily to the respiratory tract. In addition, CMA3p20 infection causes substantial damage to the mouse lung with significant inflammation, immune infiltration, and pneumonia. Importantly, mutations in CMA3p20 do not alter its ability to infect primary human cell airway cells, and the mouse adapted strain maintains similar antigenicity to wild-type SARS-CoV-2. Utilizing this model, we demonstrated infection with SARS-CoV-2 CMA3p20 provided protection from lethal challenge with the mouse-adapted SARS-CoV. Together, results indicate the utility of CMA3p20 as a model to study SARS-CoV-2 pathogenesis and immunity in standard inbred mice.

With the threat posed by SARS-CoV-2 and its emerging variants, questions have been raised on the efficacy and duration of immunity following natural infection ^19,20^. Based on the protection provided by prior SARS-CoV-2 challenge against the heterologous SARS-CoV, our results suggest immunity is more complex than neutralizing serum titer alone. While sera from mice challenged with SARS-CoV-2 CMA3p20 neutralizes SARS-CoV at ~1/150, this level of antibody does not provide sterilizing immunity in the lung. Even after SARS-CoV infection, the serum neutralization is only ~1/200 at day 7 post infection. Yet, in terms of weight loss and disease, the SARS-CoV-2 infected mice showed a significant reduction in severity and complete protection from SARS-CoV-induced lethality.

From these initial findings, the exact mechanism of protection is not yet clear. With previous studies with SARS-CoV, antibody-based immunity has led to complete protection from viral replication in the lung ^21,22^. Similar findings have been observed in animal models of SARS-CoV-2, leading to serum neutralization as a primary correlate associated with protection ^23^. However, studies with SARS-CoV have also implicated a role for cellular based immunity ^24^. Notably, initial findings have found non-neutralizing antibody responses against common cold CoV had some level of protection ^25^. Importantly, the vast majority of T-cell studies have found epitopes directed against the more conserved nucleocapsid protein ^25^. Absent in most vaccine platforms, it is unclear if the immunity stimulated by natural SARS-CoV-2 infection will mimic vaccine induced immunity and offer protection against heterologous SARS-CoV challenge.

In regard to the mouse adapted strain, CMA3p20, several of the key mutations in the spike protein have been observed in novel SARS-CoV-2 variants of concern ^26^. Starting with the spike mutations, N501Y is represented in several COVID-19 variants of concern, found in 29% of the GSAID database of reported sequences, and has been shown to improve binding to human ACE2 ^27^. In these studies, N501Y alone (CMA1) permits SARS-CoV-2 to replicate in Balb/c mice. Similarly, spike mutation K417N, also found in several variants of concern, likely augments receptor binding and drives *in vivo* disease ^28^. In contrast to the other spike mutations, H655Y is outside the RBD, has low penetrance in GSAID sequences (<0.5%), and does not have a clear functional impact ^28^. Outside the spike changes, the other mutations found in CMA3p20 are not clear in their impact. Mutations in NSP8, NSP10 (<0.5%), and E protein are rarely found in GSAID reported sequences, are not in conserved domains, and may be hitchhiking mutations ^29^. Notably, NSP6 (L37F) has been in observed in 4% of human SARS-CoV-2 isolates, suggesting possible selection ^29^. Similarly, while N change at 128 is not common ^28^, differences in disease between CMA2 and CMA1 suggesting a role in pathogenesis.

While the mouse-adapted mutations in CMA3p20 render a pathogenic virus in mice, it does not ablate the replication capacity in human cells or antigenicity relative to the wild-type SARS-CoV-2 strains. Previously described mouse adapted strains have been shown to cause significant disease but lose replication capacity in primary human airway cultures ^15^. In these studies, CMA3p20 has similar replication levels as SARS-CoV-2 WA1 in primary HAE cultures. In addition, CMA3p20-infected mice generate antibody responses capable of neutralizing wild-type SARS-CoV-2 WA1 and the mouse-adapted strain is similarly neutralized by COVID-19 sera. Importantly, passive transfer of human convalescent sera reduced disease in mice challenged with SARS-CoV-2. Together, the results indicate that CMA3p20 induces a robust immune response similar to that seen in humans and useful for understanding immunity in standard animal models.

## Methods

### Viruses and cells

The recombinant wild-type and mouse-adapted strains of SARS-CoV-2 are based on the sequence of USA-WA1/2020 isolate provided by the World Reference Center for Emerging Viruses and Arboviruses (WRCEVA) and was originally obtained from the USA Centers for Disease Control and Prevention as described ^31^. Wild-type and mutant SARS-CoV-2 as well as recombinant mouse-adapted recombinant SARS-CoV ^17^ were titrated and propagated on Vero E6 cells, grown in DMEM with 5% fetal bovine serum and 1% antibiotic/antimycotic (Gibco). Standard plaque assays were used for SARS-CoV and SARS-CoV-2 ^32,33^. All experiments involving infectious virus were conducted at the University of Texas Medical Branch (Galveston, TX) or Emory University (Atlanta, Georgia) in approved biosafety level 3 (BSL) laboratories with routine medical monitoring of staff.

### Phylogenetic tree, sequence identity heat map, and structural modeling

Spike receptor binding tables were constructed from a set of representative group 2B coronaviruses by using alignment data paired with neighbor-joining phylogenetic trees built in Geneious (v.9.1.5) using the spike amino acid sequences derived the following accession numbers: QHU79204 (SARS-CoV-2 WA1), AGZ48806 (RsSHC014), ALK02457 (WIV16), and AYV99817.1(SARS-CoV Urbani). Mouse-adapted SARS-CoV-2 structural homology models were generated using SWISS-Model ^34,35^ with the 6LZG crystal structure (RCSB Protein Data Bank) as the template structure for the spike protein and the 2AJF crystal structure (RCSB Protein Data Bank) as the template for ACE2. Homology models were visualized and manipulated in PyMOL (version 2.4.2).

### Construction of mouse-adapted mutant SARS-CoV-2

Both wild-type and mutant viruses were derived from the SARS-CoV-2 USA-WA1/2020 infectious clone as previously described ^12^. For mouse adapted virus construction, the individual mutations were synthesized and introduced into the appropriate plasmids (F1-F7) via PCR-based mutagenesis with synthesized specific primers containing corresponding mutations. The resulted plasmid was validated by further restriction enzyme digestion and Sanger sequencing. Thereafter, plasmids containing wild-type and mutant SARS-CoV-2 genome fragments were amplified and digested by restriction enzyme. The SARS-CoV-2 genome fragments were purified and ligated *in vitro* to assemble the full-length cDNA according to the procedures described previously^12,36^. *In vitro* transcription reactions then were preformed to synthesize full-length genomic RNA. To recover the viruses, the RNA transcripts were electroporated into Vero E6 cells. The medium from electroporated cells as harvested at 40 hours post transfection and served as seed stocks for amplifying one passage on Vero E6 cells (P1 stock). Viral mutants were confirmed by sequence analysis prior to use. Synthetic construction of SARS-CoV-2 mouse adapted strains were approved by the University of Texas Medical Branch Institutional Biosafety Committee.

### *In vitro* infection

Viral infections in primary human airway cells were performed as previously described ^37^. Briefly, the apical side of the HAE cultures were washed 3 times with PBS. Cultures were infected with SARS-CoV-2 WT (WA1) or CMA3p20 at MOI 0.01 and allowed to adsorb for 1 hr at 37°C. After adsorption, the apical side was washed 3 times with PBS, and the basolateral media replaced. Viral washes were collected adding PBS to the apical side and incubated for 30 minutes at 37°C. Viral titer was evaluated by focus forming assay as previously described ^37^. RNA from HAE were collected at 48 hours post infection. RNA was harvested from mock-infected and infected HAE cultures by treating with RNA lysis buffer for >5 min and gently pipetting to recover cells. Total RNA was extracted using the Zymo Quick-RNA miniprep kit (VWR; R1055) according to the manufacturer’s protocol. Purified RNA was reverse transcribed into cDNA using the high-capacity cDNA reverse transcription kit (Thermo Fisher, 43-688-13). RNA levels were quantified using the IDT Prime Time gene expression master mix and TaqMan gene expression Primer/Probe sets (IDT) and run on a QuantStudio5 qPCR system using SARS-CoV-2 RDRP-specific primers (forward [F], GTGARATGGTCATGTGTGGCGG; reverse [R], CARATGTTAAASACACTATTAGCATA) and probe (56-6-carboxyfluorescein [FAM]/CAGGTGGAA/ZEN/CCTCATCAGGAGATGC/3IABkFQ) were used. Three or more biological replicates were harvested at each described time and results are representative of multiple experiments. No blinding was used in any sample collections, nor were samples randomized. Microsoft Excel for Mac 2011 was used to analyze data.

### Deep sequencing analysis

RNA libraries of SARS-CoV-2 mutants were prepared with 300 ng of RNA using the Tiled-ClickSeq protocol as previously described ^38,39^ using tiled primers cognate to the SARS-CoV-2 genome (accession number NC_045512.2) and the TruSeq i7 LT adapter series and i5 hexamer adaptors containing a 12N unique molecular identifier (UMI). Libraries were sequenced on the Illumina MiSeq platform with MiSeq Reagent Kit v2 using paired-end reads (R1:250 cycles, R2:50 cycles). Raw data was de-multiplexed using TruSeq indexes using the MiSeq Reporter Software. Demultiplexed read data were quality filtered, adaptor-trimmed and primer-trimmed as previously described ^39^. Reads were mapped to the WA-1 reference (NC_045512.2) using *ViReMa* ^40^. Reads were de-duplicated with *umi_tools* ^41^ using 12N unique molecular identifiers (UMIs) embedded in the i5 click-adaptor. A consensus reference sequence was generated using Pilon^42^ ensuring that read coverage was greater than 25x across 99.5% of the reference genome. Pileup files were generated using Samtools v1.9 ^43^ and minority variants were extracted by nucleotide voting (PHRED>=30) using a custom python3 script previously described ^39^.

### Plaque reduction neutralization test

Neutralization assays were preformed using conventional plaque reduction neutralization assay (PRNT_50_) as previously described ^44^. Briefly, 100 PFU of SARS-CoV-2, mouse-adapted SARS-CoV-2, or SARS-CoV MA15 was incubated with serially diluted serum from mice or COVID-19 patients (total volume of 200 μl) at 37°C for 1 h. The virus-serum mixture was added to the pre-seeded Vero E6 cells. After 1 h 37°C incubation, 2 ml of 2% high gel temperature agar (SeaKem) in DMEM containing 5% FBS and 1% P/S was overlaid onto infected cells. After 2 days of incubation, 2 ml neutral red (1 g/l in PBS; Sigma) was added to the agar-covered cells. After another 5-h incubation, neutral red was removed. Plaques were counted for NT50 calculation. The PRNT assay was performed at the BSL-3 facility at UTMB.

### Ethic Statement

This study was carried out in accordance with the recommendations for care and use of animals by the Office of Laboratory Animal Welfare, National Institutes of Health. The Institutional Animal Care and Use Committee (IACUC) of University of Texas Medical Branch (UTMB) approved the animal studies under protocol 1711065 and 1707046. For samples Emory University, collection and processing were performed under approval from the University Institutional Review Board (IRB #00001080 and #00022371). Adults ≥18 years were enrolled who met eligibility criteria for SARS-CoV-2 infection (PCR or rapid antigen test confirmed by a commercially available assay) and provided informed consent.

### Human Serum samples

For Emory University, acute peripheral blood samples were collected from hospitalized patients at the time of enrollment. Infected patients were randomly selected from a convenience sample and no data was collected on the number of patients that were pre-screened or declined participation. All patients enrolled in July 2020 and had a mean age of 57 (range: 26-85; 50% male). Samples were collected in the first 9 days (range: 2-9) of their hospital stay (range: 3-33 days) and mostly 1-2 weeks after symptom onset (range 5-19 days), the majority of the patients had comorbid conditions (n=16) with 19 out of 20 having severe disease and one patient had moderate disease. All of these patients had radiological evidence of pneumonia; 19 out of the 20 patients required supplemental oxygen, and 4 out of 20 patients were admitted to the intensive care unit (ICU). Three enrolled patients died of COVID-19.

### Mice and *in vivo* infection

Ten-week-old BALB/C mice were purchased from Charles River Laboratories and were maintained in Sealsafe™ HEPA-filtered air in/out units. Prior to infection, animals were anesthetized with isoflurane and infected intranasally (IN) with 10^4^ to 10^6^ plaque forming units diluted 50 μl of phosphate-buffered saline (PBS). Infected animals were monitored for weight loss, morbidity, and clinical signs of disease, and lung titers were determined as described previously ^45^. Infected animals were weighed daily, and lung tissue collected 2-, 4- and 7-days post infection for downstream analysis by plaque assay.

### Real-Time PCR for Viral RNA

RNA from tissues were collected using RNA later (). Samples were subsequently homogenized with Trizol reagent (Invitrogen). RNA was then extracted from Triazol using the Direct-zol RNA Miniprep Plus kit (Zymo Research #R2072) per the manufacturer’s instruction. Extracted RNA was then converted to cDNA with the iScript cDNA Synthesis kit (BioRad #1708891). Quantitative real time PCR (qRT-PCR) was performed with the Luna Universal qPCR Master Mix (New England Biolabs #M3003) on a CFX Connect instrument (BioRad #1855200). Primer 1 (Forward - AAT GTT TTT CAA ACA CGT GCA G and Primer 2 (Reverse - TAC ACT ACG TGC CCG CCG AGG) were used to detect SARS-CoV-2 genomes. A primer annealing temperature of 63°C was used for all assays.

### Histological Analysis

The left lung was removed and submerged in 10% buffered formalin (Fisher) without inflation for 1 week. Lungs from mice sacrificed on day 2 and day 4 were fixed in formalin and paraffin embedded. Five-micron serial sections were taken and used for histopathological staining with hematoxylin and eosin and Immunohistochemistry (IHC) assay. IHC was conducted using rabbit polyclonal antibodies raised against the SARS-CoV nucleocapsid protein (clone NB100-56576 [1:100], Novus Biologicals, Littleton, CO) and biotinylated anti-rabbit IgG, streptavidin AP (Vector Labs, Burlingame, CA) with Permanent Red chromogen (Dako/Agilent, Santa Clara CA). Normal rabbit serum was used as primary antibody for the negative control. Briefly, the sections were deparaffinized by immersion in three xylene baths for 5 minutes each. The slides were rehydrated by immersion in a series of alcohol baths ranging from 100 to 95% for 5 minutes each. The slides were pretreated with a citrate-based buffer, pH 6.0 at 98° C for heat-induced epitope retrieval. Endogenous avidin and biotin sites were blocked using avidin/biotin blocking kit (Abcam, Cambridge, MA). The slides were incubated with anti-SARS-CoV antibody for one hour at room temperature, washed in 1X Tris-buffered saline (TBS), 0.05% Tween 20 (wash buffer), and incubated with goat biotinylated anti-rabbit IgG (H + L) antibody for 30 min at room temperature followed by rinse in wash buffer. Slides were then incubated with streptavidin-AP conjugate for 30 min at room temperature and washed, followed by incubation with Permanent Red for 5 minutes. Counterstaining with Mayer’s hematoxylin solution was performed and the slides mounted with coverslip using Permount mounting medium.

### Data Availability

The raw data that support the findings of this study are available from the corresponding author upon reasonable request.^46^

### Biological Materials

Recombinant wild-type and mutant SARS-CoV-2 described in this manuscript will be made available through the World Reference Center for Emerging Viruses and Arboviruses (WRCEVA) at UTMB through material transfer agreement. The SARS-CoV-2 B.1.1.7 strain was provided by Dr. Natalie Thornburg and colleagues at the Centers for Disease Control and Prevention.

## Acknowledgements

Research was supported by grants from NIAID of the NIH to (AI153602 and 1R21AI145400 to VDM to P-YS; R24AI120942 (WRCEVA) to SCW, P51OD011132, R56 AI147623 and U19AI090023 to MSS. AEM is supported by a Clinical and Translational Science Award NRSA (TL1) Training Core (TL1TR001440) from NIH. ALR was supported by an Institute of Human Infection and Immunity at UTMB COVID-19 Research Fund. Research was also supported by STARs Award provided by the University of Texas System to VDM, and trainee funding provided by the McLaughlin Fellowship Fund at UTMB. P-YS was also supported by CDC grant for the Western Gulf Center of Excellence for Vector-Borne Diseases, and awards from the Sealy & Smith Foundation, Kleberg Foundation, John S. Dunn Foundation, Amon G. Carter Foundation, Gilson Longenbaugh Foundation, and Summerfield Robert Foundation. MSS was also supported by the Emory Executive Vice President for Health Affairs Synergy Fund award, the Pediatric Research Alliance Center for Childhood Infections and Vaccines and Children’s Healthcare of Atlanta, COVID-Catalyst-I^3^ Funds from the Woodruff Health Sciences Center and Emory School of Medicine, Woodruff Health Sciences Center 2020 COVID-19 CURE Award.

## Author Contributions

Conceptualization, AM, MV, BAJ, ALR, MSS, XX, P-YS, VDM; Methodology, AM, MV, BAJ, MED, AV, PV, KD, ALR, DW, MSS, XX, P-YS; Investigation, AM, MV, BAJ, MED, AV, KL, CS, PV, RML, KD, ALR, DW, MSS, XX, PY-S; Resources, JAP, KSP, SCW, KD, ALR, DW, MSS, XX, P-YS, VDM; Data Curation, AM, MV, BAJ, MED, AV, PV, KD, ALR, DW, MSS, XX, PY-S, VDM.; Writing-Original Draft, AM, VDM; Writing-Review & Editing, AM, P-YS, VDM; Data Visualization, AM, KD, VDM; Supervision, SCW, ALR, DW, MSS, P-YS, VDM.; Funding Acquisition, SCW, MSS, P-YS, VDM

---

**S. Figure 1.**
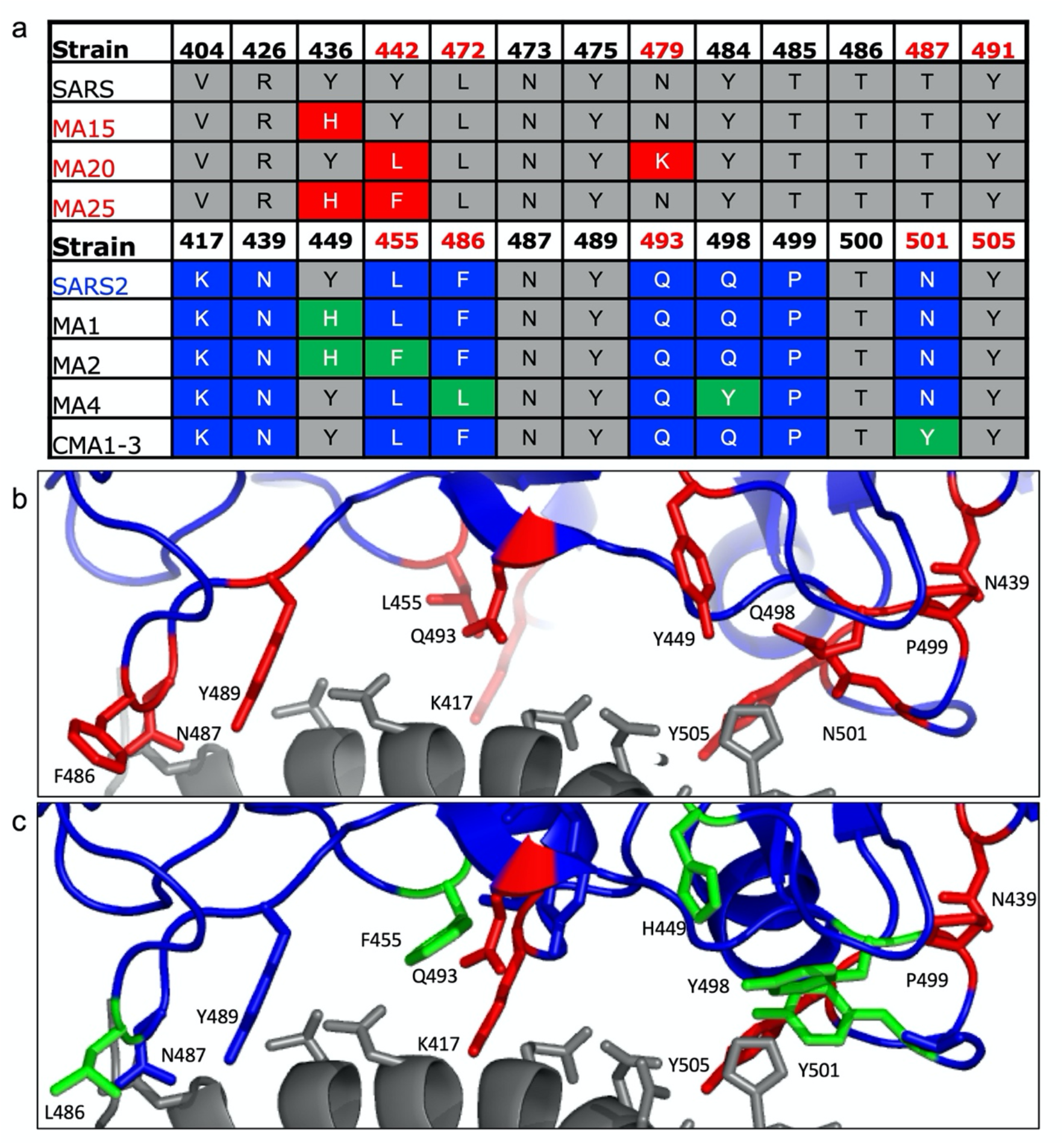
Modeling changes to mouse-adapt SARS-CoV-2. a) Key amino acid residues found in the receptor binding domain (RBD) of mouse adapted strains of SARS-CoV were aligned to SARS-CoV-2 and used to design mouse-adapted mutations ^13^. Key interaction sites between SARS-CoV spike and ACE2 molecules highlight in red ^30^. b-c) Modeling of key RBD residue interactions with mouse ACE2 (PDB:2AJF) comparing b) WT SARS-Cov-2 residues versus c) mutations (green) predicted to improve binding.

**S. Figure 2.**
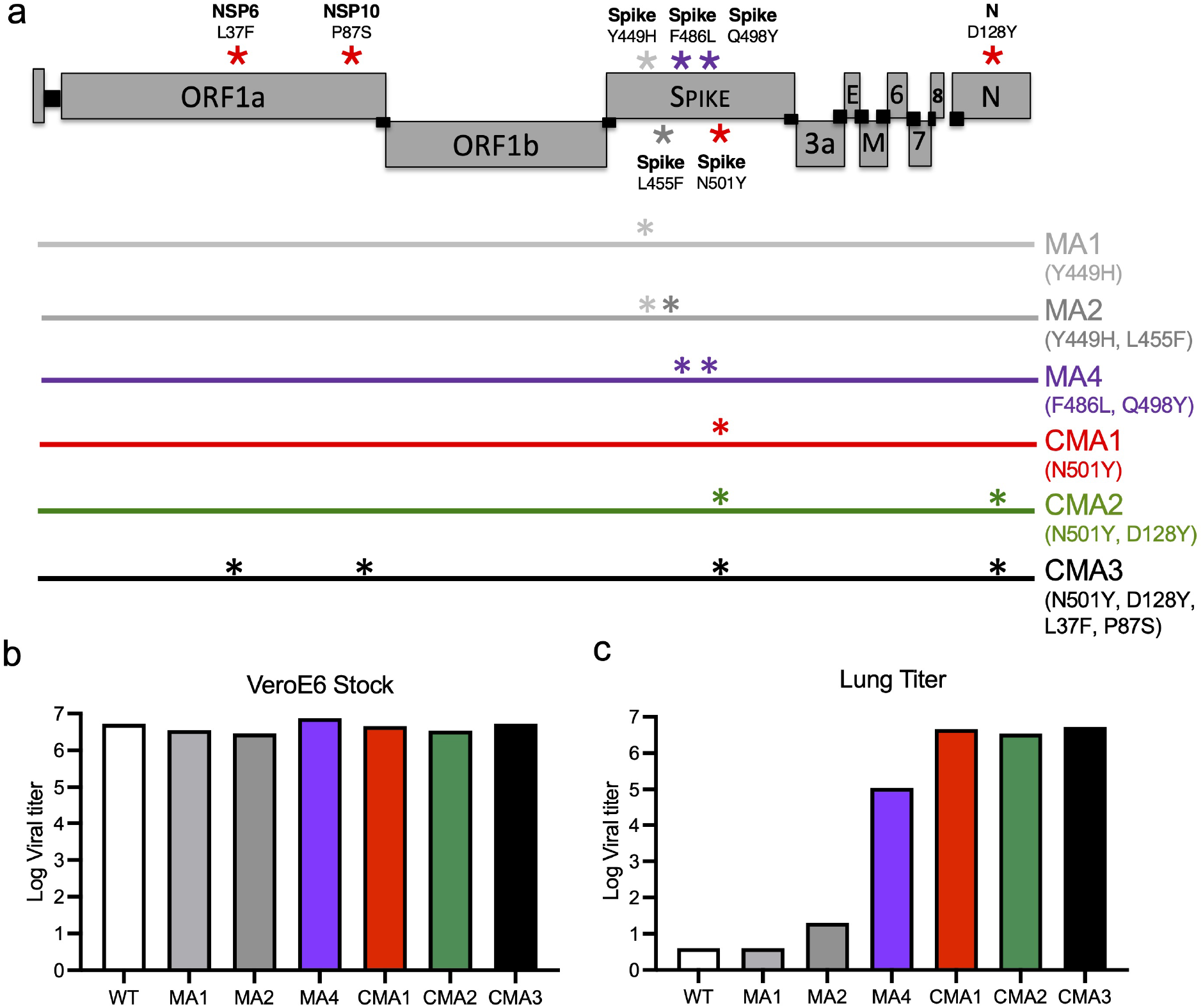
Construction of mouse-adapted SARS-CoV-2 Mutants. a) SARS-CoV-2 genome schematic indicating location of amino acid mutations for MA1, MA2, MA4, CMA1, CMA2, and CMA3. b) Viral replication of stock viruses of MA1, MA2, MA4, and CMA1-3 grown on VeroE6 cells. c) Viral replication of MA1, MA2, MA4, and CMA1-3 from lung homogenates isolated from infected mice 2 days post infection.

**S. Figure 3.**
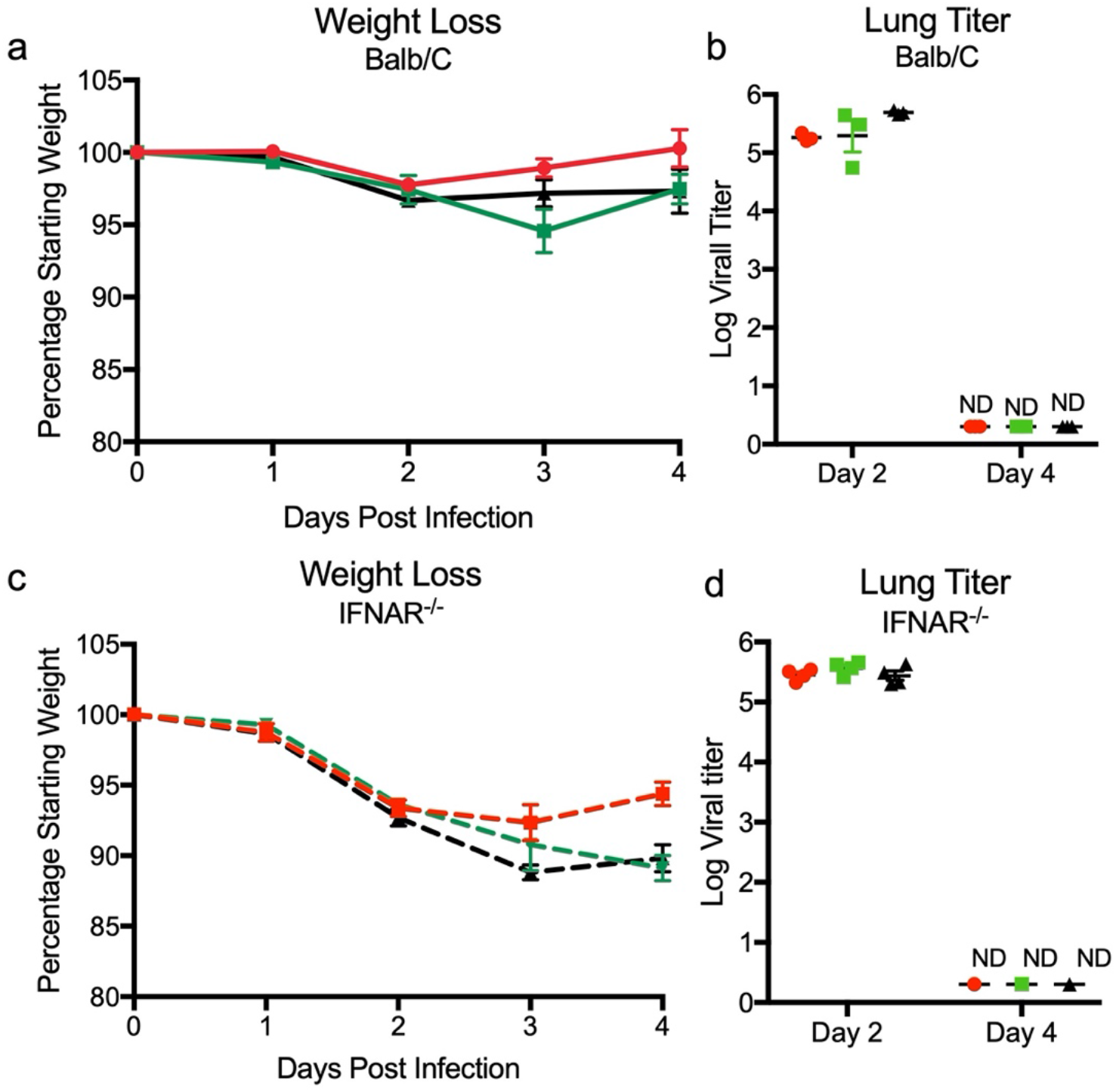
SARS-CoV-2 mutants CMA1, CMA2, and CMA3 replicate in laboratory mice. a-b) Ten-week-old female Balb/c mice infected with 10^5^ PFU of CMA1 (red), CMA2 (green), or CMA3 (black) were examined for a) weight loss and b) viral lung titer following infection at days 2 and 4. c-d) Ten- to twelve-week-old female IFNAR^-/-^ SVJ129 mice infected 10^5^ PFU of CMA1 (red), CMA2 (green), or CMA3 (black) were examined for c) weight loss and d) viral lung titer following infection at days 2 and 4. ND-non-detected.

**S. Figure 4.**
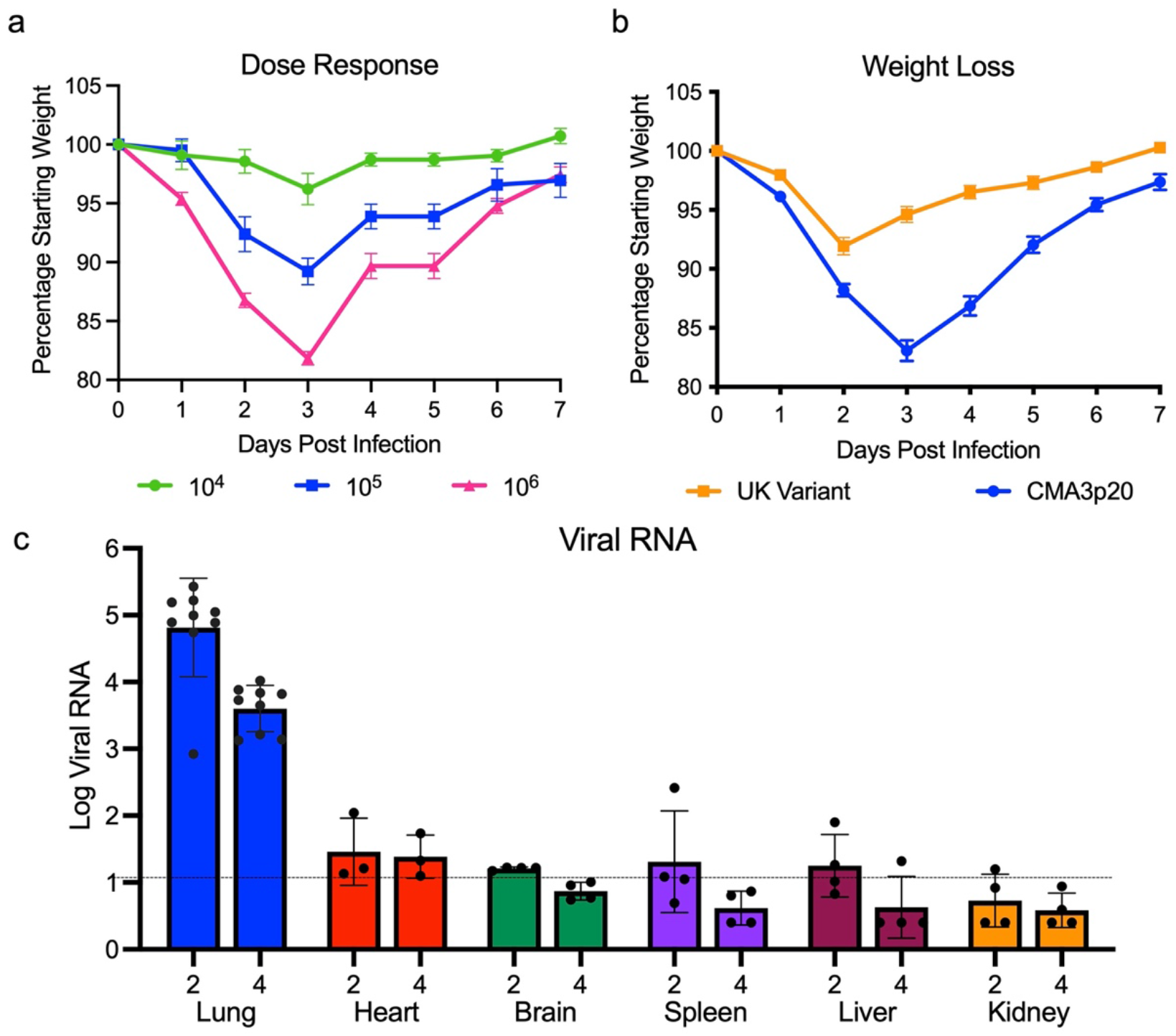
In vivo characterization of SARS-CoV-2 CMA3p20. a) Examination of ten-week-old female Balb/c mice infected with SARS-CoV-2 CMA3p20 at 10^4^, 10^5^, and10^6^ PFU (n=5). b) Comparison of weight loss in ten0-week old female Balb/c mice infected with 10^6^ PFU of SARS-CoV-2 CMA3p20 (blue) or SARS-CoV-2 variant B.1.1.7 (orange). c) RT-PCR of viral RNA load found in lung, heart, brain, spleen, liver, and kidney following 10^5^ PFU infection of SARS-CoV-2 CMA3p20 2- and 4-days post infection. Dotted line signifies viral RNA value derived from mock infected samples.

**S. Figure 5.**
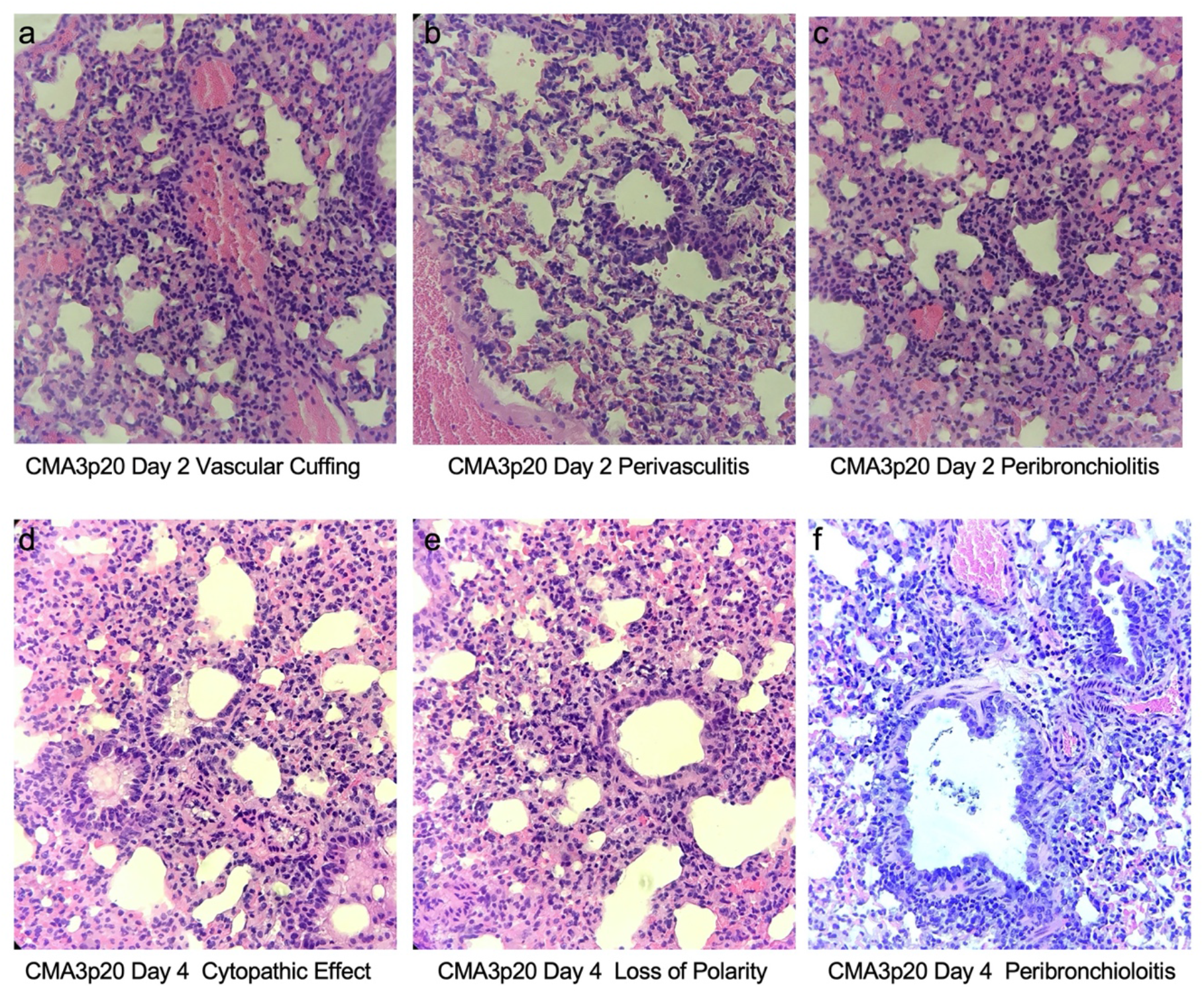
SARS-CoV-2 CMA3p20 induces significant lung damage following infection. a-c) CMA3p20 infected animals 2 days post infection showing a) perivascular cuffing, b) perivasculitis and c) peribronchiolitis. d-f) CMA3p20 induced lung inflammation and damage 4 days post infection including d) cytopathic effect of the virus, e) loss of cellular polarity, and f) inflammatory cells in the lumen. Magnification at 10x for a-f.

